# TropD-Detector: A CRISPR/LbCas12a-Based system for rapid and sensitive screening of *Trypanosoma cruzi* in Chagas vectors and reservoirs

**DOI:** 10.1101/2024.10.15.618462

**Authors:** Luis A. Ortiz-Rodríguez, Rafael Cabanzo, Jeiczon Jaimes-Dueñez, Stelia C. Mendez-Sanchez, Jonny E Duque

## Abstract

Chagas disease, also known as American Trypanosomiasis, is a zoonosis with global distribution caused by the parasite *Trypanosoma cruzi*, primarily transmitted through the feces of infected triatomines. The emergence of new cases in non-endemic areas highlights the importance of early pathogen detection in vectors and reservoirs to generate effective control strategies and establish preventive policies. The objective of this study was to design and validate a detection system of *T. cruzi* based on specific DNA cleavage and collateral activation using the CRISPR/LbCas12a system, targeting the genes Cytochrome B (*Cytb*), 18S ribosomal subunit (*SR18s*), and histone (*H2A*). This system was validated for their uses in both vectors and reservoirs of the parasite. The first step was to conduct a bioinformatic analysis of the target genes and design RNA guides for each cleavage site and primers to amplify the target region through PCR. Subsequently, we sequenced the amplified DNA target and validated the detection system using *T. cruzi* DNA extracted from naturally infected *Rhodnius pallescens* in the metropolitan area of Bucaramanga, Colombia. After standardizing the method, we tested the CRISPR/Cas system with Silvio X10/4 laboratory strain of *T. cruzi* and scaled up to blood samples of naturally infected *Didelphis marsupialis*. As a result, we observed the DNA cleavage employing the CRISPR/Cas system with *Cytb* guide with a sensitivity of up to 0.0475 ng/uL of provided DNA. Sequencing of the *Cytb* gene showed no mutations in the cleavage site. However, point mutations and indels were found in *SR18S* and *H2A*, avoiding the formation of the CRISPR/LbCas12 complex. Furthermore, we introduce the design of a fluorescent detection prototype with CRISPR/LbCas12a called “Tropical Diseases Detector” (TropD-Detector). This device operates with an excitation wavelength of 480 nm emitted by an LED and a high-pass light filter with a cutoff wavelength of 500 nm. We detected positive samples using any photographic camera system. The TropD-Detector provides a visual, viable, and sensitive method for detecting *T. cruzi* in both vectors and reservoirs from endemic areas.

## Introduction

Chagas disease (CD) is historically considered a rural and semi-rural health issue in Latin America (Duque et al., 2024). However, current reports show the spread of the disease to other continents where it was previously non-existent, positioning it as an emerging and expanding public health concern (Irish et al., 2022; Requena-Méndez et al., 2015). *Trypanosoma cruzi* is the etiological agent of CD, which affects millions of people, particularly in developing countries from America (Echeverría et al., 2020). Vectorial transmission of the parasite occurs through contact with the feces of infected insects from the Triatominae subfamily (Monteiro et al., 2018). In addition to insect transmission, other ways of infection include contaminated food ingestion, infected blood transfusion, laboratory accidents, and organ transplants have been reported. CD’s current control and prevention efforts focus on vectorial control, early disease detection, and access to treatment (Briceño-León, 2009; Duque et al., 2024). While a wide range of sensitive and specific diagnostic techniques exist in humans, in vectors and reservoirs of *T. cruzi*, the available methods for diagnosis are limited and costly, reducing the likelihood of conducting epidemiological studies in these hosts.

In vector diseases, the One Health concept comes with an integrated perspective of control that considers the interconnectedness of human, animal, and ecological health. This holistic view has gained relevance for its technical application in CD due to the significant rise in acute cases over recent decades, with more than 70% of records from oral transmission in which infected reservoirs and vectors have been involved (Shikanai-Yasuda & Carvalho, 2012). This finding suggests that current preventive strategies must expand beyond vectorial control based on documented outbreaks, even without vectorial insects (López-García & Gilabert, 2023). In this context, the reservoirs of *Didelphis* genus, take relevance due to their role in the oral transmission of *T. cruzi* through contaminated food from infected secretions of their anal glands (Ríos et al., 2011). Considering that increasing deforestation in endemic regions from Latin America has facilitated the migration of reservoirs and vectors of *T. cruzi* to urban and peri-urban areas, it is crucial to establish new systems of diagnosis and surveillance in these hosts to help at health entities to reduce the incidence of CD cases in urban centers with prevention programs.

In light of the above, to implement prevention programs, it is crucial to establish screening and diagnostic systems targeting species involved in the parasite’s life cycle across all geographical regions affected by CD (Olivera et al., 2024; Velásquez-Ortiz et al., 2022). In this way, early detection of the parasite in vectors or reservoirs can identify active transmission cycles and pinpoint high-risk areas for communities, thus enabling the implementation of localized preventive measures (Cantillo-Barraza et al., 2015; Velásquez-Ortiz et al., 2022). Although there is significant progress in the detection systems for *T. cruzi*, it is essential to highlight that most current methods primarily focus on humans. Moreover, depending on each technique, concerns regarding sensitivity, reliability, and high costs exist. For instance, in Latin America, the most simple and rapid detection test is the peripheral blood smear, which can be a viable initial screening tool with an effectiveness of 68% as long as it is applied in patients with the acute phase of the disease (Duque et al., 2024). More complex techniques include serological tests, which have higher detection capacity and are common in hospitals. Nonetheless, false positives with CD have been reported with other parasites responsible for leishmaniasis, especially in geographic regions endemic to both diseases (Abras et al., 2016; Gomes et al., 2009). In addition to these techniques, it is necessary to employ confirmatory diagnostic essays such as indirect immunofluorescence (IFI) or molecular tests.

Among molecular techniques, endpoint PCR is considered the “Gold Standard” test for molecular identification of *T. cruzi*. While this method is highly reliable, it is prone to false positives due to cross-contamination between samples. Its effectiveness depends on the appropriate primer design and the operator’s technical expertise (Stadhouders et al., 2010). Despite the diagnostic system’s advantages in CD, false negatives could also appear in host low parasitic loads (Hirschhorn et al., 2023; Viljoen et al., 2005). More sensitive techniques, such as real-time PCR (qPCR) or digital PCR (dPCR), have been conducted in various diseases, including CD. Given its high sensitivity, qPCR is an alternative to PCR in chronic cases, which tends to produce false negatives in such instances (Ramírez et al., 2018; Seiringer et al., 2017). However, the downside of these methods lies in the need for robust, expensive equipment with highly trained personnel and having limited field applicability. Additionally, it is necessary to bring the samples to a third-level laboratory, which results in extended detection times to get a diagnosis.

Given the context mentioned above, it is essential to develop new diagnostic techniques that are accurate, confirmatory, rapid, cost-effective, and adaptable to settings with limited infrastructure, such as rural areas in Latin America where the CD is endemic (Alarcón de Noya & Jackson, 2020). Addressing these diagnostic necessities before explained, CRISPR/LbCas12a systems adapted as detection sensors are a new, sensible, low-cost, programmable, and specific alternative to detect target regions within DNA. Although CRISPR systems were originally a genome editing tool, their application in disease diagnosis has increased in recent years, mainly due to their role in rapid COVID-19 tests during the pandemic (Y. Chen et al., 2022). The CRISPR-based detection methodology involves a few steps. Initially, a previous amplification of the target site, designed as a guide-RNA, is necessary (Kaminski et al., 2021). Conventional techniques, such as PCR or isothermal amplification methods like RPA or LAMP, achieve this pre-amplification (Li et al., 2019). Subsequently, Cas protein and designed guide-RNA are complexed to locate the target site. Once identified, the Cas protein produces the first (*Cis)* cleavage at a specific site on the amplified DNA. Consecutively, a second (*Trans*) random cleavage occurs, which cleavages a single-strand DNA fluorescent reporter (East-Seletsky et al., 2016). It is excited about a wavelength of 480 nm; the reporter emits detectable fluorescence at 530 nm. Within the visible spectrum, this fluorescence is in the blue light. It is employed to develop pathogen detection systems that use blue LEDs or robust equipment as a UV light transilluminator (Bai et al., 2022; Ding et al., 2020; Qian et al., 2021; Zhou et al., 2022).

In light of the previously discussed challenges in diagnosis like costs and the need for a simple screening method, the objective of this study is to develop a detection prototype of *T. cruzi* employing CRISPR/LbCas12a system, applicable to insect vectors and parasite reservoirs to establish a highly specific, sensitive, rapid, programmable, cost-effective, and portable detection system that any person with a short training can use for the detection of *T. cruzi*.

## Methods

### Bioinformatic analysis

To achieve molecular detection of *T. cruzi*, we targeted conserved DNA regions to enable specific detection of the parasite. A thorough literature review led to selection of three genes: SR*18S, H2A*, and *Cytb* (Brisse *et al*., 2001, 2003; Pavia *et al*., 2007). Representative sequences from the NCBI database of each gene were downloaded and aligned using the SNAPGENE software. Consequently, we designed primers for the *SR18s* and *H2A* gene, while primers for the *Cytb* gene were adapted from previously reported for *T. cruzi* detection (Brisse et al., 2003). We designed guide RNAs (gRNAs) for each gene with 24-pb length and a PAM sequence TTTV in the CHOPCHOP v2 software (J. S. Chen et al., 2018). These gRNAs were chosen based on high specificity for *T. cruzi*, efficiency scores, and low off-target effects in other genome regions. Then, we filtered the designs in BLAST and selected the most optimal for each gene. In like manner, designed primers were analyzed in BLAST to select the most specific design for *T. cruzi*. Each primer was compared and aligned against negative controls with *Rhodnius prolixus, Leishmania infantum, Trypanosoma theileri, and Lutzomyia* sp (**Figure 1 A)**.

**Figure 1.**
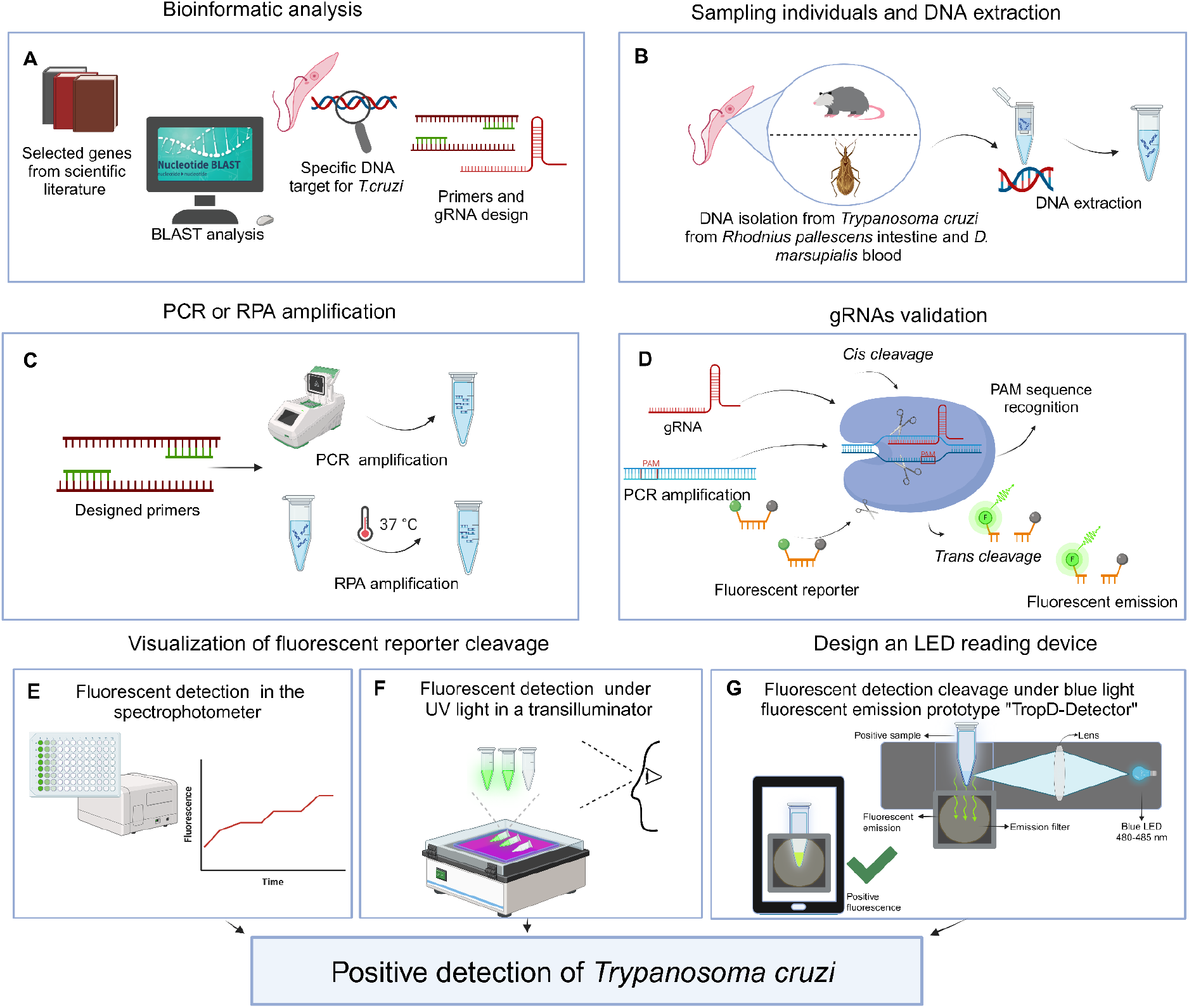
Schematic of the CRISPR/LbCas12a assay workflow for detection of *T. cruzi* in reservoirs and vectors of the parasite. **A. Bioinformatic analysis**. BLAST analysis and literature review of target regions from *T. cruzi*. **B. Sampling individuals and DNA extraction**. Isolation DNA of *T. cruzi* from collected samples from *D. marsupialis* and intestine of *R. pallescens*. **C. PCR or RPA amplification**. PCR amplification in thermocycler or isothermal amplification with RPA. **D.gRNAs validation**. In amplified DNA, the gRNA guides the LbCas12a protein cleavage (Cis cleavage). Then, the protein cleaves the fluorescent single-stranded DNA reporter due to its Trans activity. The reporter is excited at 480 nm and emits fluorescence at 530 nm **E. Fluorescent detection in the spectrophotometer**. The emitted fluorescence is measured in a spectrophotometer over two hours at 37 °C. **F. Fluorescent detection under UV light in a transilluminator**. Visualization of positive samples under UV light on a transilluminator. **G. Fluorescent detection cleavage under blue light fluorescent emission prototype “TropD-Detector**. Employing blue LED light, the reporter is excited, and the emitted fluorescence is registered and detected in a conventional cellphone. Figure created with BioRender.com.

### DNA isolation from *T. cruzi* from *R. pallescens* intestine

The collection of *R. pallescens* specimens was conducted between November 2021 and February 2022 by personnel from the Medical Entomology Laboratory in local neighborhoods of Bucaramanga, Colombia. The sampling was carried out with Angulo trap (Angulo & Esteban, 2011), in three neighborhoods with forest fragments. A total of 26 specimens were collected: twelve from the “Pan de Azúcar” neighborhood, eight from “Limoncito,” and six from “Bucarica.” DNA extraction we made with Qiagen DNeasy Blood and Tissue protocol (Qiagen, Valencia, CA, USA) from dissected upper intestinal samples and stored at -20°C.

### Blood sample collection and DNA extraction of *D. marsupialis*

Collected samples from *D. marsupialis* were obtained from specimens treated at the Wildlife Rescue and Care Center of the metropolitan area of Bucaramanga. Initially, these animals were weighed employing a digital scale with a 20 kg capacity (Vibra, Terrace, USA®). A veterinarian anesthetized each specimen using xylazine (doses of 2.2 mg/kg) and ketamine hydrochloride (doses of 5-7.5 mg/kg). Ultimately, 0.5mL - 1 mL blood samples were isolated from the caudal vein for each animal, employing 3 mL syringes and 23G x 1” needles, and stored in microtubes with EDTA. DNA extraction from *D. marsupialis* blood samples was conducted with the Corpogen extraction Kit, following the manufacturer’s protocol (CorpoGen, Bogotá, Colombia®). Briefly, 250 μL of blood with EDTA was extracted, with the final elution carried out in 100 μL of elution buffer. Finally, the quality and quantity of the extracted DNA were assessed using a NanoDrop 2000 (Thermo Fisher Scientific, Massachusetts, USA).

For wild insects, Decree 1376 of June 27, 2013, was taken into account, which regulates the collection permit for specimens of wild species of biological diversity for non-commercial scientific research purposes. The Medical Entomology Laboratory, part of CINTROP-UIS, operates under Framework Permit for Specimen Collection No. IDB0398, issued by the National Authority of Environmental Licenses. The collection of *R. pallescens* was conducted following laboratory animal handling protocols and approved by the Scientific Research Ethics Committee (CEINCI) Comité de ética en investigación científica CEINCI” of the Industrial University of Santander (acronym in Spanish CEINCI), as recorded in Minutes No. 15, dated August 27, 2021. Nucleotides used in this study were sourced from the project “Producción de nucleótidos a partir de biomasa residual de la agroindustria para diagnóstico por biología molecular en el departamento de Santander,” also approved by CEINCI under Minutes No. 20, dated May 20, 2022.

### PCR amplification

Designed primers were validated through PCR using DNA isolated from the intestine of *R. pallescens* infected with *T. cruzi*. We employed the Q5® Hot Start High-Fidelity 2× Master Mix (New England Biolabs, American), in a T100 thermocycler (Bio-RAD Inc). After validation, we amplified DNA samples of *T. cruzi* from the Silvio strain culture and blood samples of *D. marsupialis* naturally infected with the parasite (**Figure 1 B)**. The amplified products were visualized on a 2% agarose gel stained with SYBR® Safe DNA Gel Stain (Invitrogen, #S33102). Electrophoresis was performed at 100 volts and 90 milliamps for 50 minutes. Designed primers were also tested against various DNA samples used as negative controls, including *Rhodnius prolixus, Leishmania infantum, Trypanosoma theileri*, and *Lutzomyia* sp (**Annex Figure 1**).

### gRNAs validation

Designed gRNAs were synthesized commercially by IDT (Integrated DNA Technologies, Inc.) and validated through *in vitro* cleavage of PCR products, which were observed on agarose gel using amplified *T. cruzi* DNA extracted from *R. pallescens*. We adapted the methodology from *the in vitro* digestion protocol by New England Biolabs (M0653), with concentrations modified during the standardization. CRISPR reactions were performed in a 30 μL reaction mixture consisting of 6 μL of 300 nM LbCAS12a protein, 6 μL of 300 nM gRNAs, and 6 μL of NEB reaction buffer, incubated at 37° C for ten minutes. To visualize the DNA cleavage, 15 μL of the reaction mix was run on a 2% agarose gel stained with SYBR® Safe DNA Gel Stain (Invitrogen, #S33102).

### Target gene sequencing

After gRNA validation, the DNA amplified was sequenced from *T. cruzi* genes *SR18s, H2A*, and *Cytb* extracted from *R. pallescens* intestine. Bidirectional sequencing was conducted through Macrogen’s sequencing service. The obtained sequences were aligned and compared with reference sequences obtained from the NCBI database. The results were analyzed using the SnapGene software.

### Visualization of fluorescent reporter cleavage

The fluorescent system methodology was based on J. S Chen et al. 2018. We selected a single-stranded DNA (ssDNA) reporter probe labeled with a fluorophore and quencher (FQ) (56-FAM/TTATT/3IABkFQ), which was synthesized commercially by IDT. (J. S. Chen et al., 2018). The ssDNA reporter, gRNA, and LbCas12a protein concentrations were adapted as previously described (Lee et al., 2020). The fluorescence reaction mixture was adjusted to 100 nm of ssDNA reporter, while the final concentration of gRNA and LbCas12 was set at 60 nm. An equivalent volume of protein was used in the NEB buffer; molecular water was added to achieve a final volume of 50 μL.

We quantified the emitted fluorescence using dark plates in a SYNERGY H1 spectrophotometer, taking measurements every three minutes at 37° C for two hours, with 40-second shaking intervals between readings. The samples were subsequently visualized on a UV transilluminator, allowing visual identification of positive samples. To evaluate the sensitivity of the CRISPR/LbCas12a system, we diluted the amplified DNA to final concentrations of 40 ng/uL, 20 ng/uL, 10 ng/uL, 5 ng/uL, 1 ng/uL. The same approach was applied to assess the sensitivity of PCR combined with CRISPR/LbCas12 system by performing serial dilutions from 1:2 to 1: 32 using *T. cruzi* DNA extracted from cultures of the parasite with an initial concentration of 9.5 ng/uL. The DNA diluted was amplified through PCR using the *Cytb* gene and observed on a 2% agarose gel. Cleavage by the CRISPR system was then evaluated using the amplified DNA from these dilutions with the designed guide. Each assay was performed in triplicate.

### Design an LED reading device

We designed a portable device using affordable materials. It features a darkened cavity to prevent external light. At one end, a blue LED emits light in the range of 480-485 nm is powered by a conventional 9-volt battery. A lens with a focal length of f=5 cm focused the light onto a PCR tube sample. The emitted fluorescence from the sample is observed at a 90° angle relative to the incident light. A high-pass filter was used with a cutoff wavelength of 500 nm. A camera was positioned behind the filter to capture the image of the excited sample. A light trap was positioned adjacent to the sample to reduce the intensity of reflected light. Fluorescence visualization of the samples was achieved using a standard mobile phone. We validated the samples obtained from the culture, insects, and reservoir. Subsequently, we confirmed the system’s sensitivity using samples derived from the dilution of the amplified product and the dilution of the DNA provided to the PCR. All the assays were compared with the results obtained under UV light visualization and in the spectrophotometer quantification.

### Statistical analysis

Fluorescence readings from the spectrophotometer were normalized as reported (Dueñas et al., 2022). We first employed a Kolmorgorov-Smirnov test to evaluate the normality of the replicates. Kruskal-Wallis test was used to evaluate differences in non-normal data, followed by a post hoc Dunn’s test, while an one-way ANOVA was conducted in normal data, followed by a Tukey post hoc test. The homogeneity of the replicas was applied to the normalized data in the sensitivity tests (**Annex Table 2)**. All analyses were executed using Statistica Version 10.

## Results

### Bioinformatic analysis

Selected primers from bioinformatic analysis produced a PCR product length between 600 pb and 900 pb. BLAST analysis revealed 100% homology with *T. cruzi* for the designed primers. The three analyzed genes were tested on species used as negative controls, as shown in **Annex Table 1**. Ten possible designs were obtained within the amplified region for each gene, with potential binding sites containing PAM (TTTV) motifs. One gRNA was selected for each gene, prioritizing high specificity and 0% off-target score, as shown in **Table 1**.

**Table 1.**
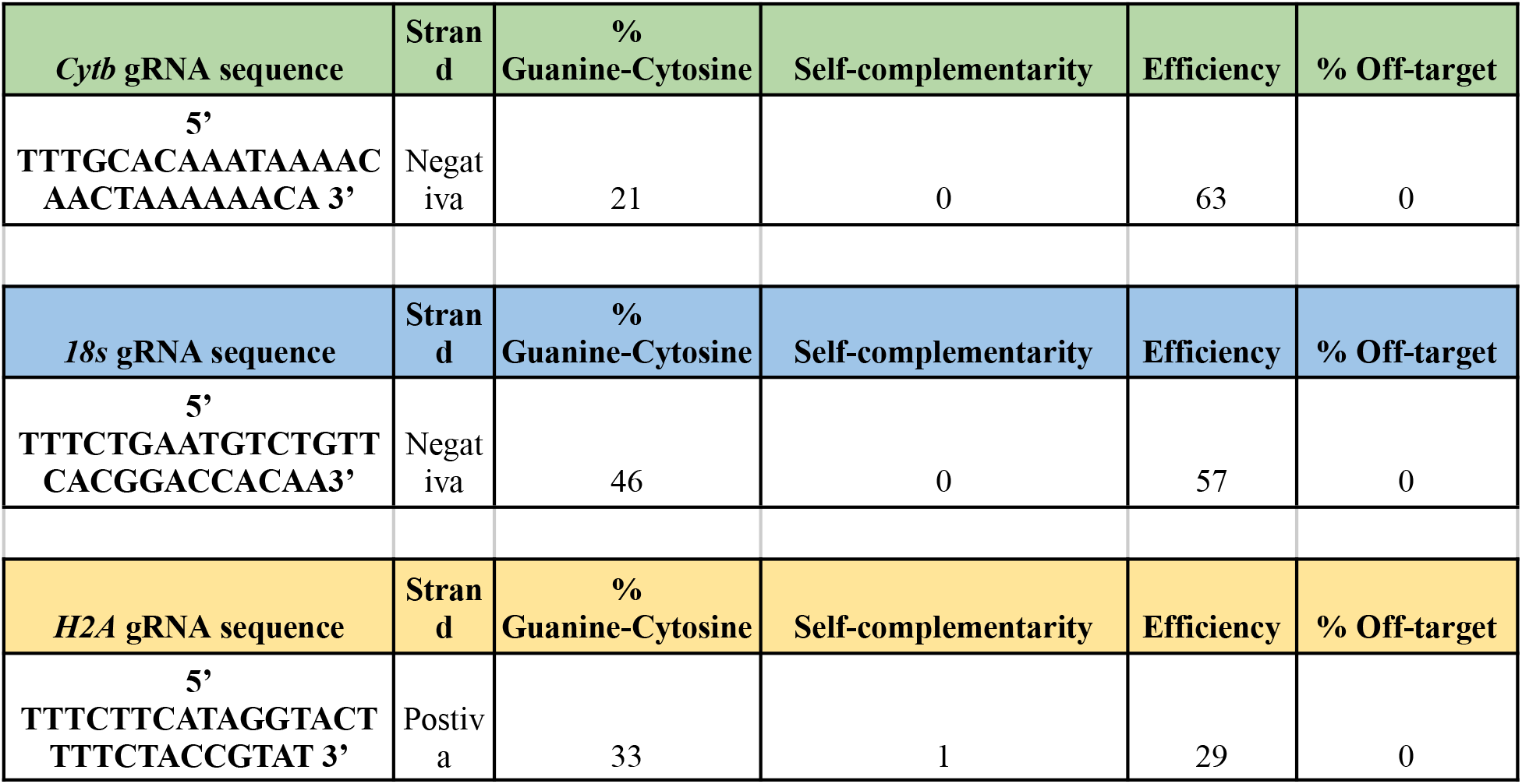
gRNAs designed for each gene. The “strand” column indicates the orientation of the guide RNA; if labeled “Negative,” it refers to the complementary strand. The “% Guanine-Cytosine “ reflects the proportion of G and C within the guide sequence. “Self-complementarity” represents the probability of the gRNA binding to itself. “Efficiency” refers to the likelihood of the sequence binding with the target site. The “% Off-target” represents the percentage of possible non-target sites within the total DNA of the species where the gRNA might bind.

### PCR amplification

We amplified the *Cytb, SR18s*, and *H2A* genes in different samples by PCR, including negative controls (see **Annex 1)**. The PCR product was positive in the three *T. cruzi* samples. In *T. cruzi* DNA extracted from the intestines of *R. pallescens*, a faint band was observed at a concentration of 18 ng/μL, whereas a strong band was observed in *T. cruzi* from culture at 9.5 ng/uL. Unexpectedly, positive bands were also observed in the negative controls. For example, a weak band was detected in *T. theileri* at 203 ng/uL for both the *SR18s* and *H2A genes*. Similarly, in *L. infantum* at 9.9 ng/uL, positive amplification was observed for the *SR18s* gene, while ambiguous amplification occurred for the *Cytb* gene. In *Lutzomya* sp, at 3.9 ng/uL, a weak band was detected for the *SR18s* gene and ambiguous amplification for the *H2A* gene. No amplification was observed in the DNA from *R. pallescens* at 23.9 ng/μL.

### gRNAs validation

The designed primers successfully amplified the target site for cleavage by CRISPR/LbCas12a system in the three target genes. Using the *Cytb* gRNA, two distinct bands were observed on an agarose gel, corresponding to the fragments generated at the cleavage site of the guide-protein-DNA complex. The larger fragment corresponded to a 481 pb band, while the shorter fragment measured 241 pb **(Figure 2E)**. However, no cleavage was detected for the *SR18S* and *H2A* gRNAs, as assessed by agarose gel visualization and fluorescent emission. In contrast, the gRNA yielded fluorescence levels exceeding 20,000 relative fluorescence units (RFU), as shown in **Figure 2D**. The gRNA was validated using *T. cruzi* DNA extracted cultures, as well as samples from the intestine of *R. pallescens*, and blood samples of the opossum *D. marsupailis* **(Figure 2F)**.

**Figure 2.**
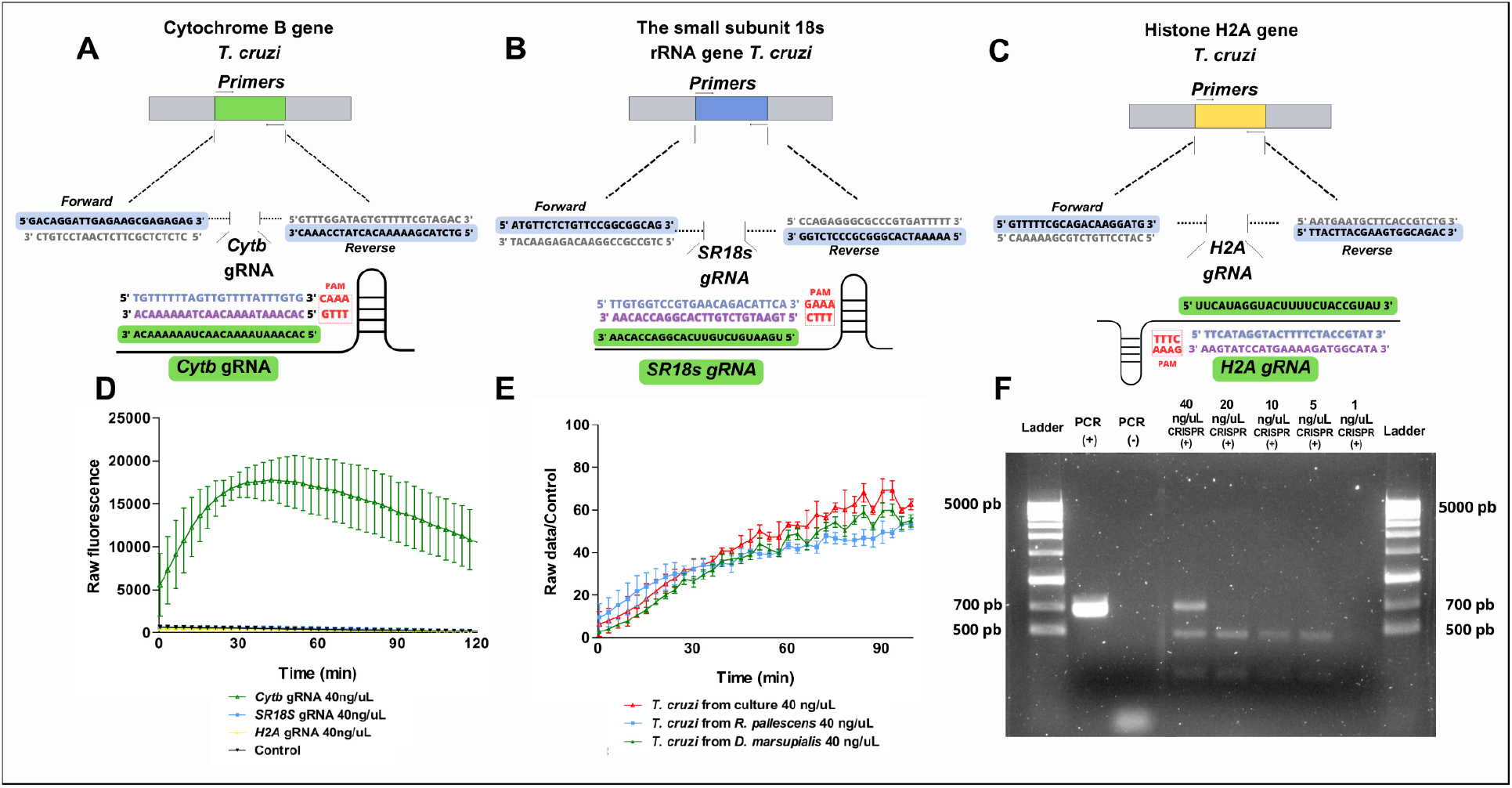
**A-C**. Representation of the *Cytb, SR18S*, and *H2A* genes selected for the identification of *T. cruzi*, along with specific primers and gRNAs design. **D**. Fluorescence intensity graph displaying raw data for each designed gRNA. No DNA was added to the Control. The *Cytb* gRNA showed significant fluorescence with p<0.05 **F**. Cleavage evaluation of diluted amplified products. **E**. Comparative graph of normalized fluorescence (Raw data/Control) corresponding to three *T. cruzi* samples. The culture sample demonstrated a significant difference from *T. cruzi* extracted from the vector’s intestine (p=0,000189) and the reservoir (p=0,024).

The amplified *Cytb* gene product was diluted to concentrations of 40 ng/uL, 20 ng/uL, 10 ng/uL, 5 ng/uL, and one ng/uL, and subsequently cleaved using the CRISPR/LbCas12a system with the corresponding gRNA. Cleavage was observed at all dilutions, as evidenced by agarose gel visualization **(Figure 2E)**. At the 1 ng/uL concentration, a weak band corresponds to the larger fragment of 481 pb length.

### DNA Sequencing

Sequencing of *T. cruzi* samples extracted from the vector’s intestine revealed point mutations in the gRNA binding region of the *SR18S* and *H2A* genes. In the *SR18S* gRNA binding site, a double insertion was detected near the PAM site. Similarly, in the *H2A* gRNA binding region, two base mutations and one indel were detected. In contrast, no variations were found in the *Cytb* gRNA binding site **(Figure 3)**.

**Figure 3.**
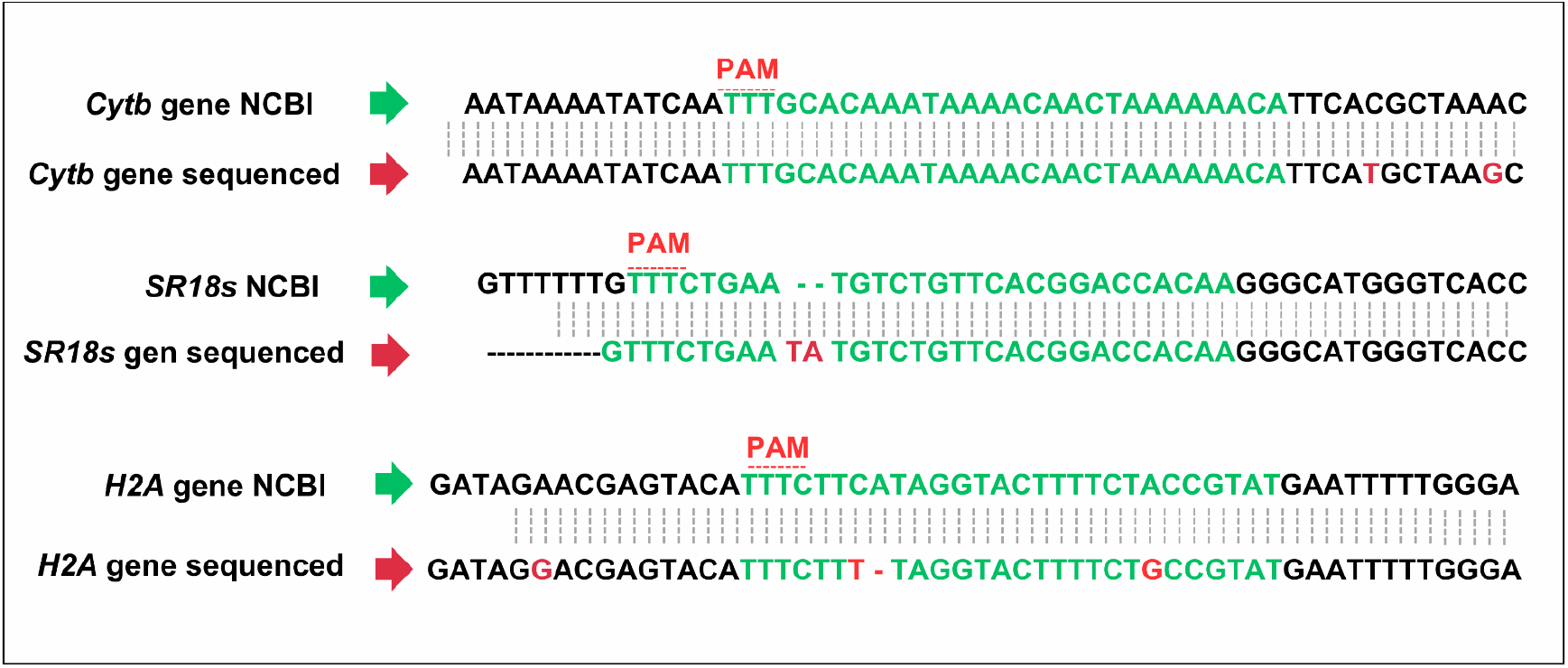
Sequencing of the target site from amplified *T. cruzi* DNA extracted from the intestine of *Rhodnius pallescens* in Bucaramanga, Colombia. The green bases indicate the gRNA binding site for each gRNA target gene. The PAM site consists of TTTV nucleotides. All results can be observed in **Annex Figure 3**.

### Amplified DNA diluted cleavage

The CRISPR/LbCas12a system’s cleavage efficiency was assessed using diluted *T. cruzi* amplified DNA from culture by measuring fluorescence emission in a spectrophotometer. We observed fluorescence emission at all dilution levels for both the cultured DNA **(Figure 4A)** and extracted from the vector’s intestine **(Figure 4C)**. The results from both sample types were normalized by dividing the fluorescence values for each measurement by the corresponding control without DNA **(Figure 4 B-C)**. Additionally, fluorescence emission increased proportionally with the amplified DNA. The homogeneity of replicates for each dilution was confirmed with a Kruskal-Wallis test (*p> 0*.*05*). A transilluminator evaluated the fluorescence mix in PCR tubes under UV light. We observed fluorescence in all dilutions with a green color, clearly distinguishable from the control without DNA (**Figure 4 E-F)**.

**Figure 4.**
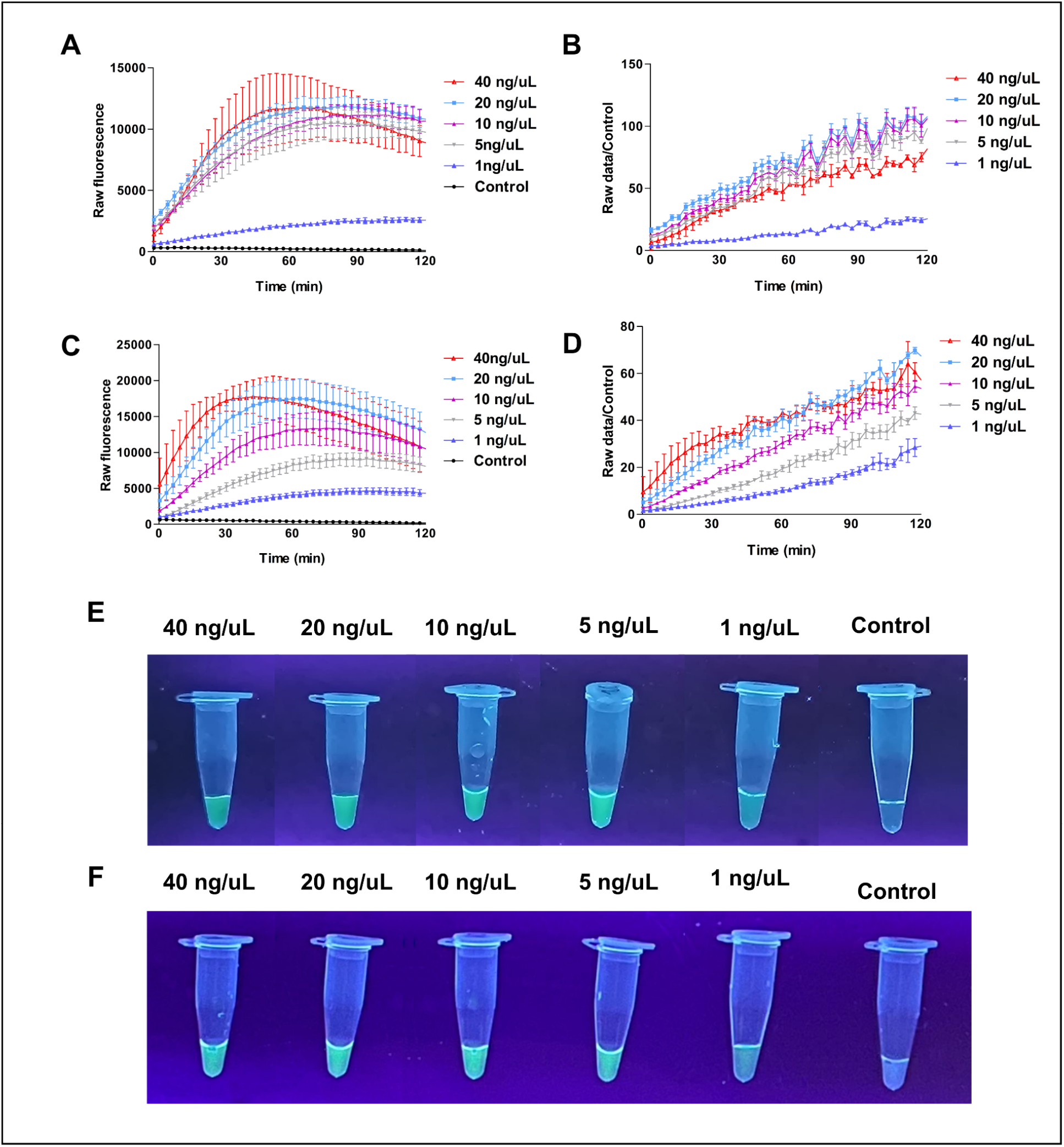
**A**. Raw fluorescence values from diluted *T. cruzi* DNA amplified *Cytb* gene from culture, cleaved by the CRISPR/LbCas12a system. **B**. Normalized fluorescence data emitted by the cleavage of diluted amplified DNA from culture. **C**. Raw fluorescence data from CRISPR/LbCas12a cleaving in the *Cytb* gene of *T. cruzi* amplified DNA diluted from the intestine of *R. pallescens*. **D**. Normalized fluorescence data emitted by the cleavage of diluted amplified DNA extracted from the intestine of *R. pallescens* **E-F**. Visual cleavage detection of *T. cruzi* from culture and extracted from the intestine of *R. pallescens* under UV light using a transilluminator.

### Fluorescence emission analysis using the “TropD-Detector” device

The designed device enabled the fluorescence excitation of the ssDNA reporter using LED light powered by a 9-volt battery. This simple and practical device facilitated qualitative evaluation of diluted samples, avoiding the use of carcinogenic UV radiation, unlike the transilluminator. Fluorescence was recorded using a conventional smartphone in a darkened room (**Figure 5A)**. The emission was properly filtered, producing a green color in the PCR tubes containing positive samples. *T. cruzi* samples derived from the vector’s intestine and reservoirs exhibited fluorescence at 40 ng/uL, clearly differentiable from the control **(Figure 5 B)**. In the dilutions of amplified DNA, visible fluorescence was observed at all evaluated dilutions, except for the control tube without DNA (**Figure 5C**).

**Figure 5.**
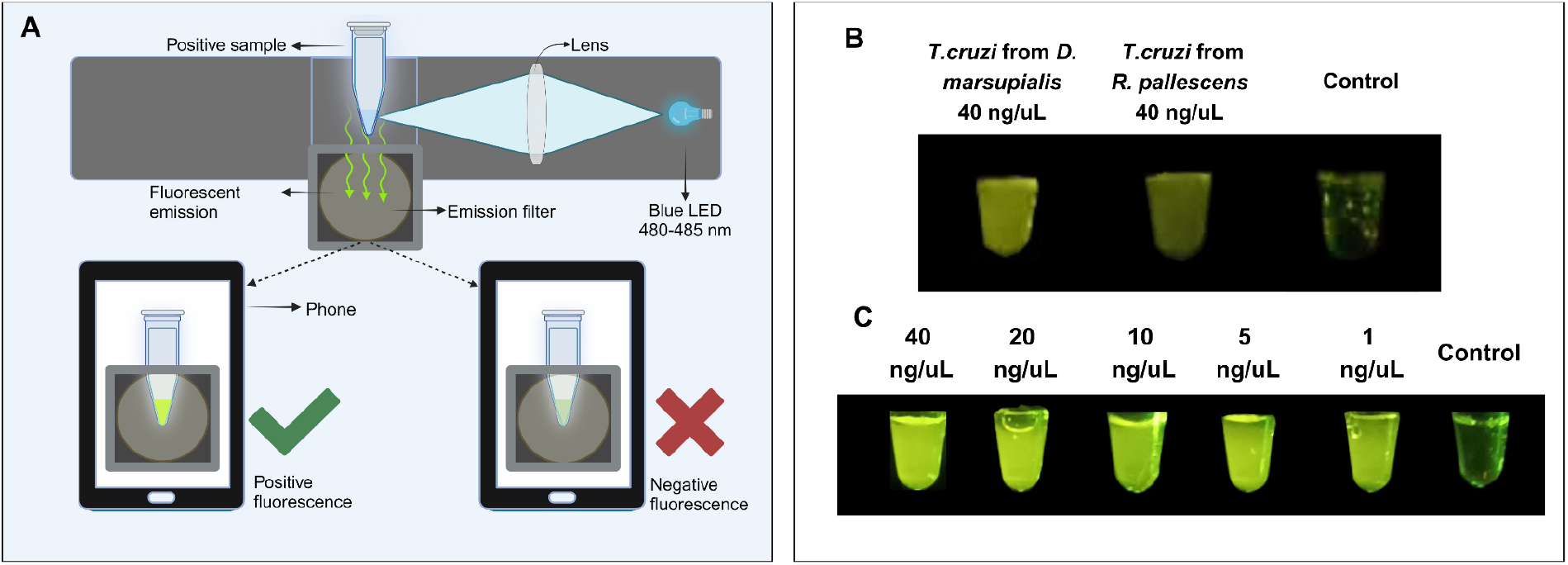
TropD-Detector device. **A**. The fluorescent emission prototype utilizes blue light at 480 nm to excite the sample containing the CRISPR/LbCas12a system and the ssDNA fluorescence reporter. The emitted fluorescence passes through a light filter, enabling visual detection using a smartphone. Positive samples display a bright green color, while the negative control exhibits a green color with a transparent background. **B**. Fluorescence emission from positive samples of *T. cruzi* extracted from *D. marsupialis* and the intestine of *R. pallescens* **C**. Visual cleavage detection of diluted DNA amplified from *T. cruzi* culture. Figure created with BioRender.com.

### Analytical Sensitivity testing with pre-amplified DNA dilution

The analytical sensitivity of the CRISPR/LbCas12a system was evaluated by diluting the DNA sample prior to PCR. We realized serial dilutions ranging from 1:2 to 1:32 using *T. cruzi* DNA derived from culture, starting with an initial concentration of 9.5 ng/uL. Positive bands were observed down to a concentration of 0.0475 ng/uL (1/16 dilution), with band intensity decreasing proportionally to the DNA concentration **(Figure 6A)**. Cleavage results were consistent across all visualization systems with the PCR amplifications. No amplification was detected at the 1/32 dilution, which aligned with the absence of fluorescence in both the spectrophotometer and the UV transilluminator **(Figures 6B-D)**, as well as with the results from the TropD-Detector device **(Figure 6E)**. The fluorescence data emitted by the dilution were similar to those of the control without DNA; however, significant differences were observed (Kruskal-Wallis test *p < 0*.*05*)).

**Figure 6.**
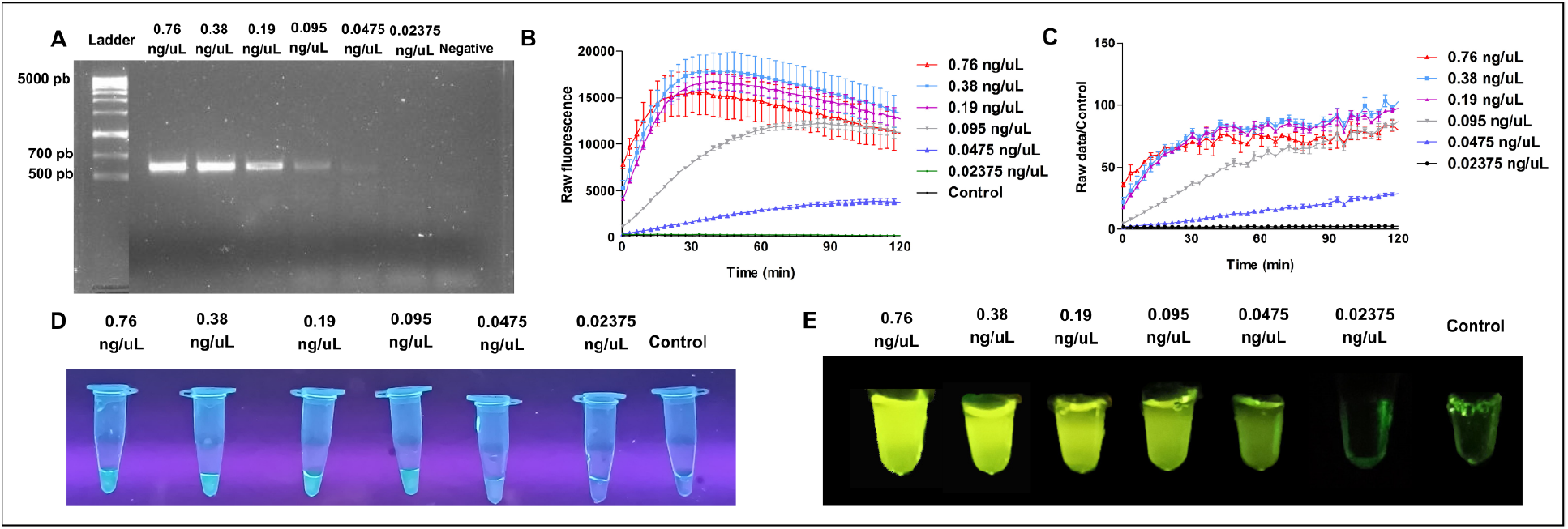
**A**. Diluted *T. cruzi* DNA from the culture before PCR, starting at 0.76 ng/uL, with no band detected at the 0.02375 ng/uL dilution. **B**. Raw fluorescence data from CRISPR/LbCas12a cleavage in the *Cytb* gene of amplified products of the pre-PCR dilutions obtained from culture. **C**. Normalized fluorescence data emitted by the cleavage of amplified products of the pre-PCR dilutions obtained from culture. **D**. Visual cleavage detection of diluted DNA under UV light using a transilluminator. **E**. Visual cleavage detection of diluted DNA prior to PCR from *T. cruzi* culture in the TropD-Detector device.

## Discussion

The CRISPR/LbCas12a system, employed in this study in conjunction with the TropD-Detector device, enabled the detection of *T. cruzi* in samples obtained from vectors and reservoirs. Its versatility allowed the development of a portable device capable of identifying positive samples using a smartphone or other image-capturing devices. As mentioned before, the positive PCR amplification observed in samples designated as negative control highlights the potential for false-positive findings in molecular diagnostics (Seiringer et al., 2017; Stadhouders et al., 2010). The methods used in this study validate the CRISPR technology as a viable alternative for parasite detection, complementing traditional amplification techniques such as PCR in sample screening.

Although the gRNAs were designed through bioinformatic analysis of various available sequences in the *SR18s* and *H2A* genes, we observed indels and point mutations in the target sites that impeded efficient binding. These differences were located within 5 to 10 bases of the PAM site, a critical area known as the seed region in Cas proteins (Moon & Liu, 2023). Previous studies have shown that mismatches in this sensitive region reduce the gRNA’s recognition capacity, as reported for Cas12 (Mann & Pitts, 2022). In contrast, no mutations were found between the designed gRNA and the target site in the *Cytb* gene **(Figure 2)**, as confirmed by the fluorescence emitted using the *Cytb* gRNA **(Figure 3 D)**, demonstrating efficient binding.

We reduced the ssDNA reporter to a final concentration of 100 nM, optimizing the number of reactions while lowering the cost per sample. This concentration aligns with the findings of Lei (2022) in detecting *Toxoplasma gondii*. It is half of the concentration reported by Dueñas (2022) for *Leishmania* sp., and is ten times lower than that mentioned by Lee 2020 in *Plasmodium* sp (Dueñas et al., 2022; Lee et al., 2020; Lei et al., 2022).

Sensitivity testing with pre-amplified dilution enabled detection up to 0.0475 ng/uL. Various factors, such as the amplification technique, sample type, or targeted gene, can influence sensitivity. For instance, the CRISPR/Cas detection system combined with RPA amplification has been applied in zoonotic diseases caused by parasites such as human African trypanosomiasis, toxoplasmosis, or malaria, achieving a detection sensitivity as low as 0.03 parasites/μL for the *SR18s* gene, as reported in two species of *Plasmodium* sp (Lee et al., 2020; Lei et al., 2022; Sima et al., 2022).

The results obtained using the TropD-Detector device were consistent with those from UV transilluminator and spectrophotometer quantification, showing a decreased fluorescence intensity with DNA decreasing concentration. The device demonstrated greater sensitivity in differentiating the 1 ng/uL concentration from the control without DNA compared to transilluminator images. This improved sensitivity is attributed to the high-pass light filter, which effectively prevents interference from the blue LED light with the fluorescence intensity emitted by the positive samples. The CRISPR-Cas system for detection using blue light for reporter excitation has been applied in previous studies. For instance, Yu et al. (2021) utilized this system to detect *Cryptosporidium parvum*, the parasitic agent of a disease known as cryptosporidiosis, primarily affecting the digestive system in humans. However, this method typically requires specialized equipment for sample visualization, such as the ‘Tanon 5200 Fluorescence Imager” or a portable gel imagining system (Yu et al., 2021). Other researchers have employed this methodology by developing specially designed cells for sample positioning (Fozouni et al., 2021), which is unnecessary for the TropD-Detector device, as the samples can be placed in standard PCR tubes.

Developing a molecular identification system requires extensive bioinformatics analysis that focuses on identifying highly conserved regions of the pathogen while accounting for potential mutations in different geographical areas, which could lead to false-negative results. These intra-species variations must be considered before field implementation. They can be addressed through prior characterization in the study area or by designing multiple guides targeting different sequences within the selected region to ensure reliable pathogen detection. We present a device suitable for low-infrastructure settings, with the potential to be scalable for use with human patients suspected of having CD.

## Conclusion

The TropD-Detector device successfully detected *T. cruzi* in samples from vectors, reservoirs, and cultures of the parasite, demonstrating the adaptability and scalability of the CRISPR/Cas system for portable, precise, practical, and cost-effective methodologies. These features are critical for molecular diagnostic tests, particularly in facilitating patient screening for tropical diseases.

## Acknowledgments

To Gustavo Adolfo Rincón and Juliana Cuadros Martínez for their collaboration in the collection of biological material and identification of the triatomines.

## CRediT authorship contribution statement

**Luis A. Ortiz-Rodríguez:** Performed experimental assays, formal analysis, investigation, methodology, TropD-Detector device designing and writing original draft, Writing review & editing. **Rafael Cabanzo:** TropD-Detector device designing, writing – review & editing. **Jeiczon Jaimes-Dueñez:** Supplying DNA samples from *D. marsupialis*, writing – review & editing. **Stelia C. Mendez-Sanchez:** Project administration, resources, writing – review & editing. **Jonny E Duque** Project design, formal analysis, investigation, project administration, resources, supervision, writing – original draft, writing – review & editing. All authors contributed and approved the manuscript.

