## Supplemental material for "TropD-Detector: A CRISPR/LbCas12a-Based system for rapid and sensitive screening of *Trypanosoma cruzi* in Chagas vectors and reservoirs"

#### Annex table 1

| | **Analysis of species in BLAST with *Cytb* primers** | | | | | |
|---|---|---|---|---|---|---|
| | **Forward** | | | **Reverse** | | |
| | 5’ GACAGGATTGAGAAGCGAGAGAG ‘3 | | | 5’ CAAACCTATCACAAAAAGCATCTG 3’ | | |
| **Specie** | **% Coverage** | **% Identity** | **Number of shared bp** | **% Coverage** | **% Identity** | **Number of shared bp** |
| *Trypanosoma theileri* | 100 | 100 | 23 | 100 | 100 | 24 |
| *Leishmania infantum* | 100 | 100 | 23 | 100 | 94 | 24 |
| *Rhodnius pallescens* | 69 | 100 | 9 | 41 | 100 | 9 |
| *Lutzomyia* sp. | 69 | 100 | 15 | 100 | 100 | 17 |

| | **Analysis of species in BLAST with *SR18s* primers** | |
|---|---|---|
| | **Forward** | **Reverse** |
| | 5’ ATGTTCTCTGTTCCGGCGGCAG ‘3 | 5’ AAAAATCACGGGCGCCCTCTGG 3’ |

| **Specie** | **% Coverage** | **% Identity** | **Number of shared bp** | **% Coverage** | **% Identity** | **Number of shared bp** |
|---|---|---|---|---|---|---|
| *Trypanosoma theileri* | | | | | | |
| *Leishmania infantum* | 95 | 100 | 15 | 100 | 100 | 15 |
| *Rhodnius pallescens* | 54 | 100 | 9 | 72 | 100 | 8 |
| *Lutzomya* sp. | 100 | 100 | 15 | 100 | 100 | 14 |

| | **Analysis of species in BLAST with H2A primers** | | | | | |
|---|---|---|---|---|---|---|
| | **Forward** | | | **Reverse** | | |
| | 5’ GTTTTTCGCAGACAAGGATG ‘3 | | | 5’ TTACTTACGAAGTGGCAGAC ‘3 | | |
| **Specie** | **% Coverage** | **% Identity** | **Number of shared bp** | **% Coverage** | **% Identity** | **Number of shared bp** |
| *Trypanosoma theileri* | 60 | 100 | 12 | 85 | 100 | 17 |
| *Leishmania infantum* | 100 | 100 | 14 | 90 | 100 | 14 |
| *Rhodnius pallescens* | 40 | 100 | 8 | 35 | 100 | 7 |
| *Lutzomyia* sp. | 100 | 100 | 14 | 100 | 100 | 14 |

.

### Annex Figure 1

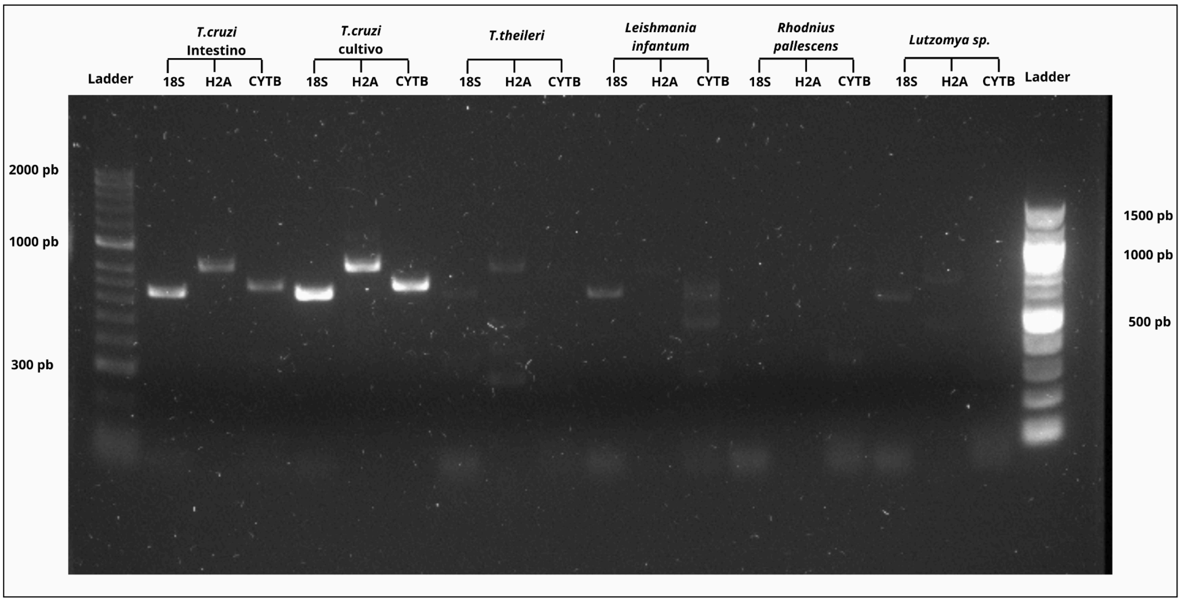


**Annex 1.** Validation of primers for the *Cytb, 18s,* and *H2A* genes in DNA from *T. cruzi* Silvio strain*, T. cruzi* extracted from the intestine of *Rhodnius pallescens, Trypanosoma theileri, Leishmania infantum, Rhodnius pallescens, and Lutzomyia* sp.

### Annex Table 2. Statistics of the graphs

| Figure | Description | Details |
|---|---|---|
| **Figure 2E** | Homogeneity of guide RNA (gRNA) replicas for *SR18s*, *H2A*, and *Cytb* | Replicas are homogeneous with *p > 0.05* |
| **Figure 2F** | Homogeneity of DNA extracted from culture, *Rhodnius pallescens* intestine, and reservoir samples | Culture: *p = 0.343*, Vector: *p = 0.4160*, Reservoir: *p = 0.5182* |
| **Figure 4B** | Normalized fluorescence data emitted by the cleavage of diluted amplified DNA from culture | 40 ng/μL: *p = 0.4164*, 20 ng/μL: *p = 0.2404*, 10 ng/μL: *p = 0.240*, 5 ng/μL: *p = 0.2130*, 1 ng/μL: *p = 0.1531*, Control: *p = 0.1343* |
| **Figure 4D** | Normalized fluorescence data emitted by the cleavage of diluted amplified DNA extracted from the intestine of *R. pallescens* | 40 ng/μL: *p = 0.4164*, 20 ng/μL: *p = 0.2404*, 10 ng/μL: *p = 0.240*, 5 ng/μL: *p = 0.2130*, 1 ng/μL: *p = 0.1531*, Control: *p = 0.1343* |
| **Figure 6C** | Normalized fluorescence data graph for pre-PCR dilutions | 0.76 ng/μL: *p = 0.2055*, 0.38 ng/μL: *p = 0.0821*, 0.19 ng/μL: *p = 0.4785*, 0.095 ng/μL: *p = 0.3633*, 0.0475 ng/μL: *p = 0.4570*, 0.02375 ng/μL: *p = 0.5073* |

### Annex  Figure 3. Sequencing Results

#### *Cytb* FORWARD

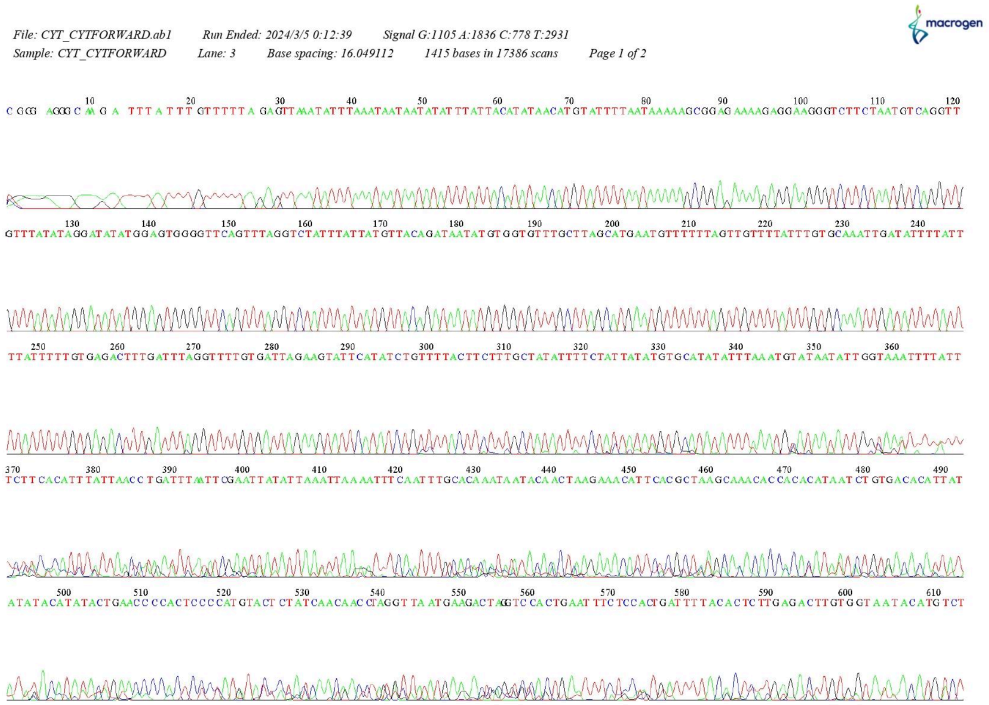

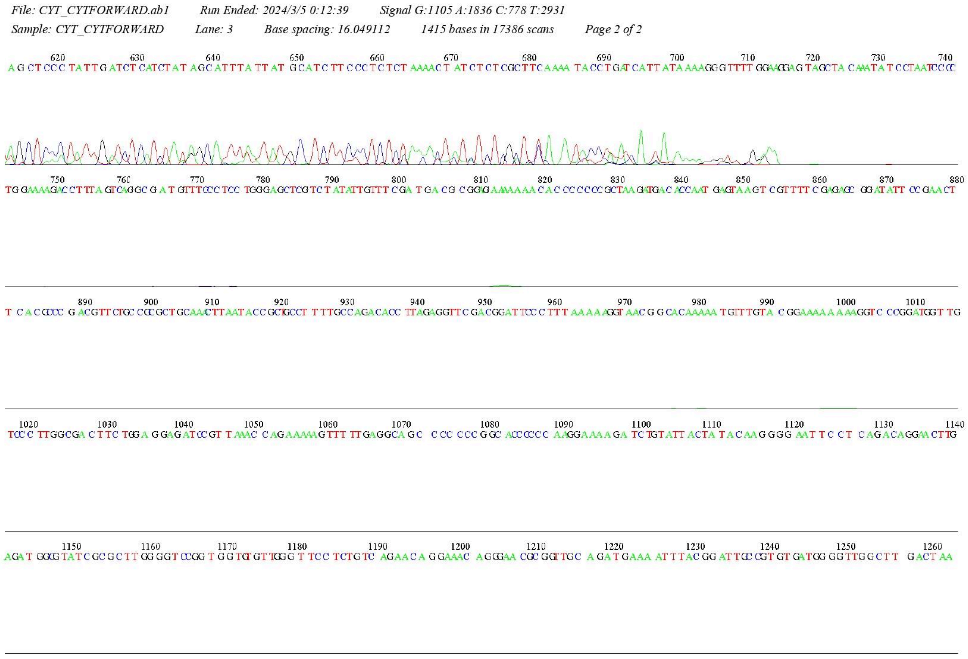


#### *Cytb* REVERSE

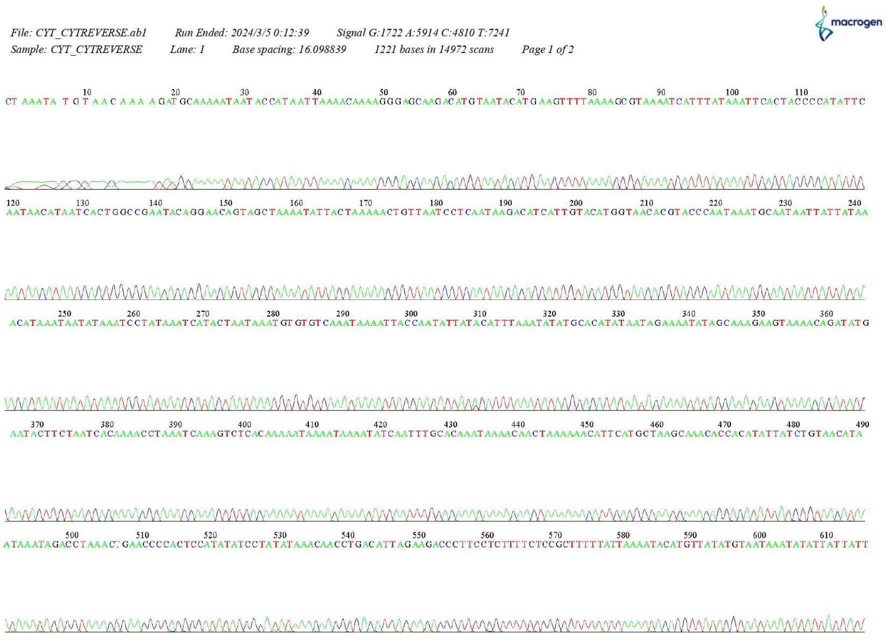

File: CYT_CYTREVERSE.ab1 Run Ended: 2024/3/5 0:12:39 Signal G:1722 A:5914 C:4810 T:7241
Sample: CYT_CYTREVERSE Lane: 1 Base spacing: 16.098839 1221 bases in 14972 scans Page 2 of 2

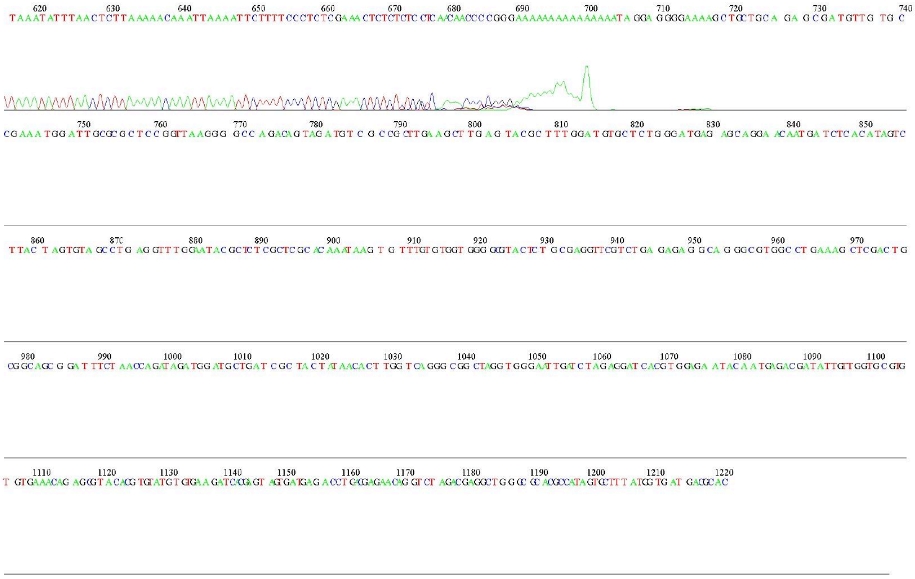


### SR*18s* FORWARD

File: 18S_18FORWARD.ab1 Run Ended: 2024/2/9 0:58:36 Signal G:300 A:445 C:577 T:591
Sample: 18S_18FORWARD Lane: 52 Base spacing: 15.765961 1761 bases in 20998 scans Page 1 of 2

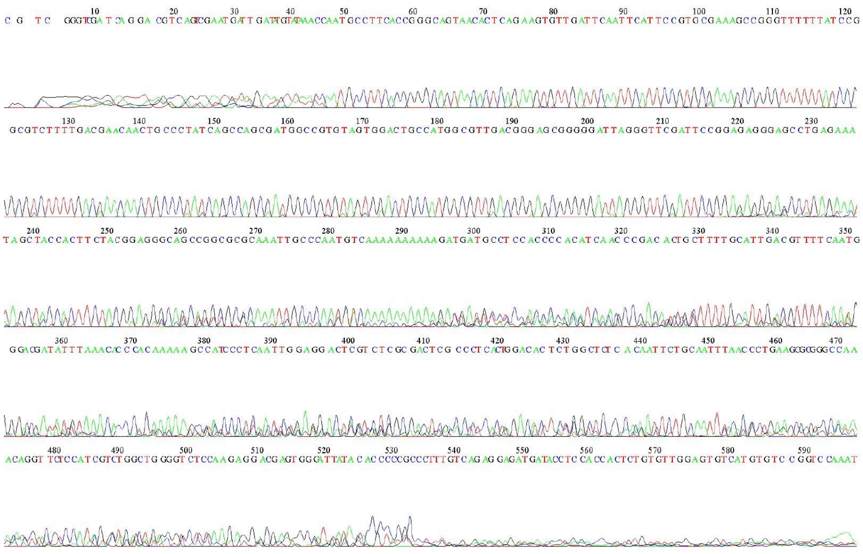

File: 18S_18FORWARD.ab1 Run Ended: 2024/2/9 0:58:36 Signal G:300 A:445 C:577 T:591
Sample: 18S_18FORWARD Lane: 52 Base spacing: 15.765961 1761 bases in 20998 scans Page 2 of 2

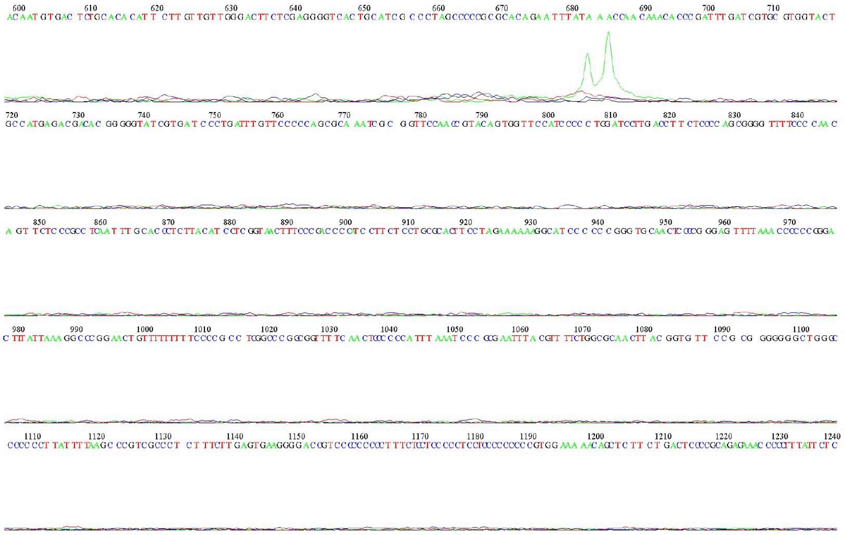

### SR*18s* REVERSE

File: 18S_18-REVERSE.ab1 Run Ended: 2024/2/9 0:58:36 Signal G:1717 A:2418 C:3458 T:3165
Sample: 18S_18-REVERSE Lane: 50 Base spacing: 16.078657 1440 bases in 18045 scans Page 1 of 2

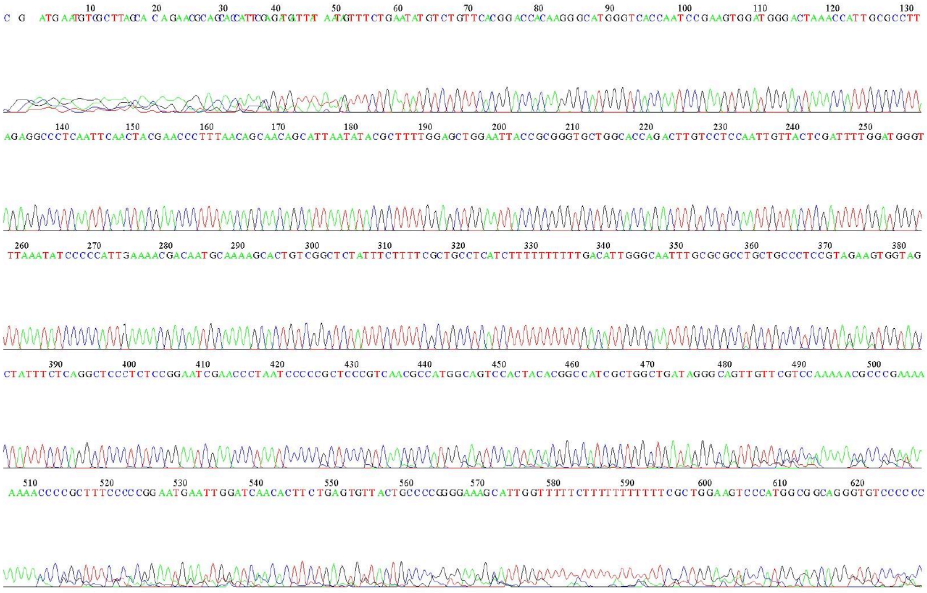

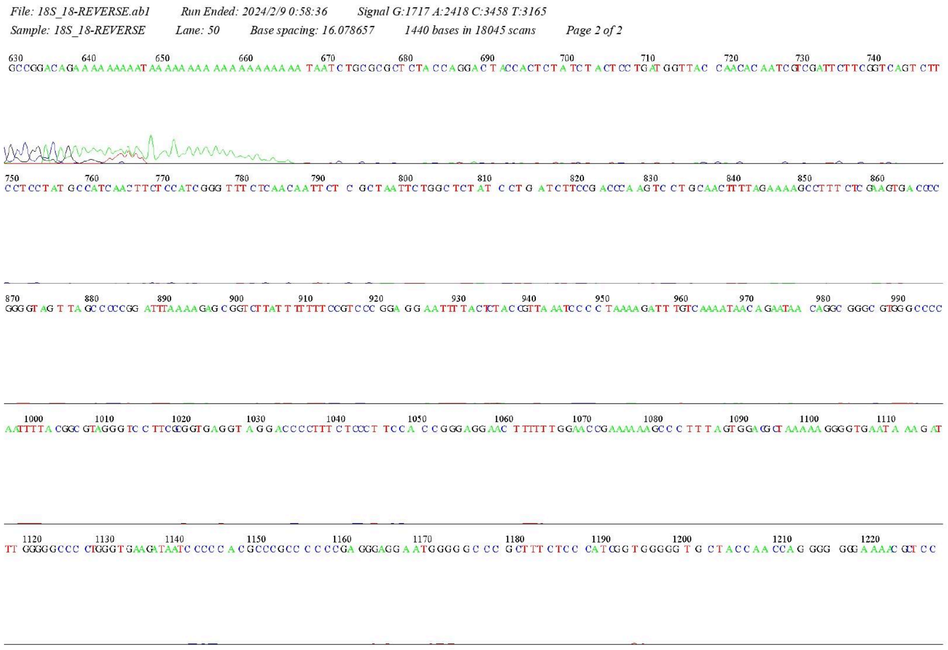
File: 18S_18-REVERSE.ab1 Run Ended: 2024/2/9 0:58:36 Signal G:1717 A:2418 C:3458 T:3165
Sample: 18S_18-REVERSE Lane: 50 Base spacing: 16.078657 1440 bases in 18045 scans Page 2 of 2

#### *H2A* FORWARD

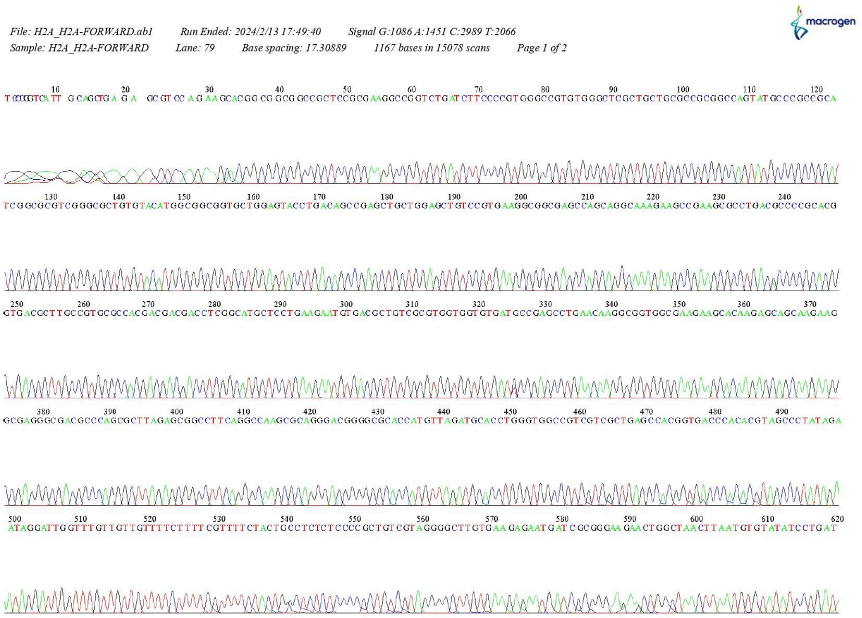
macrogen
File: H2A_H2A-FORWARD.ab1 Run Ended: 2024/2/13 17:49:40 Signal G:1086 A:1451 C:2989 T:2066
Sample: H2A_H2A-FORWARD Lane: 79 Base spacing: 17.30889 1167 bases in 15078 scans Page 1 of 2

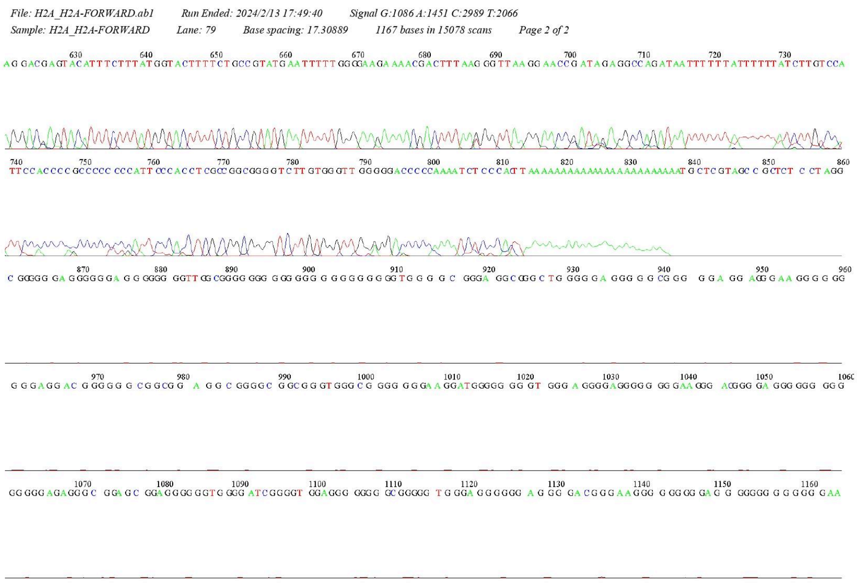
File: H2A_H2A-FORWARD.ab1 Run Ended: 2024/2/13 17:49:40 Signal G:1086 A:1451 C:2989 T:2066
Sample: H2A_H2A-FORWARD Lane: 79 Base spacing: 17.30889 1167 bases in 15078 scans Page 2 of 2

### *H2A* REVERSE

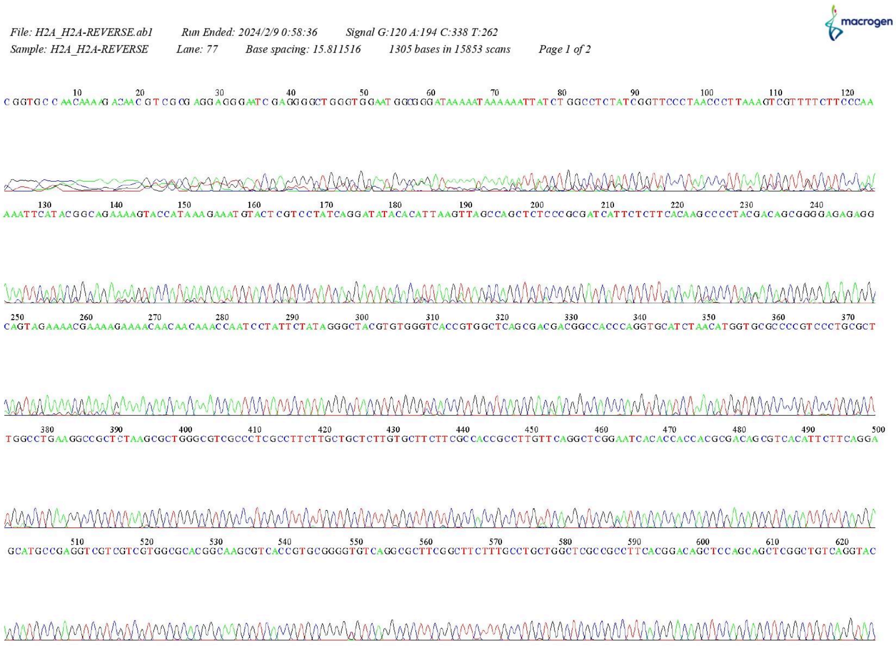
File: H2A_H2A-REVERSE.ab1 Run Ended: 2024/2/9 0:58:36 Signal G:120 A:194 C:338 T:262
Sample: H2A_H2A-REVERSE Lane: 77 Base spacing: 15.811516 1305 bases in 15853 scans Page 1 of 2

*File: H2A_H2A-REVERSE.ab1*    *Run Ended: 2024/2/9 0:58:36*    *Signal G:120 A:194 C:338 T:262*

*Sample: H2A_H2A-REVERSE*    *Lane: 77*    *Base spacing: 15.811516*    *1305 bases in 15853 scans*    *Page 2 of 2*

630 640 650 660 670 680 690 700 710 720 730 740 750

TCCAGCACCGCCGCCATGTACACAGCGCCCGACGCGCCGATGCGGCGGGCATACTGGCCGCGGCGCAGCAGCGAGCCCACACGGCCCACGGGGAAGATCAGACCGGCCTTCGCGGAGCGGCCGCC

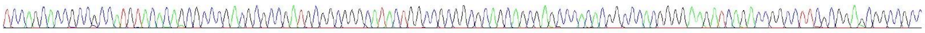

760 770 780 790 800 810 820 830 840 850 860 870

GCCGTGCTTCTTGGACGCCTTCTTCGCTGCCTGCTTCGGGGTTGCCATCCTTGTCT G CCGAAAAACAAAAA AAAAAAT CTTCAAAACTC CT ACC CTC TTCGCTAC ACGGT GAGAGGCCGG

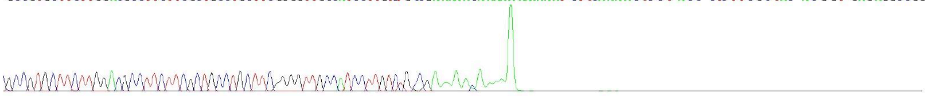

880 890 900 910 920 930 940 950 960 970 980

AT AG GGAGGG TTTG GA CGG C T TTGA T AGGT TT A TCGCC GGAATCA AATTGGGGGT CCC TGCTTCCTC GC GGGGC GGAT GGCCCCCGGCC CGCGGCGTTGGAAGGAGAGGA GGTTG

990 1000 1010 1020 1030 1040 1050 1060 1070 1080 1090 1100

AATCGT GCGGACGACTTGGG GCGAA AGCGAAGG CAACGATGCGGGGAAGGGA GG AG G GGGAGCCCGAGCCT GTTTGT TAAGG GGGGGA TA ATCAAT GGGCTTG GGGGC TGG AGTGGG G GGTT

1110 1120 1130 1140 1150 1160 1170 1180 1190 1200 1210 1220 1230

GGTT AGGTAGGCGTGGTT GAGGCGGG GAGATTGGT AGA GTGGTGGGTG AACGGCTG CCCTTGGCGTCGCGGGGA T TATTGGATG TGG GG GTGAGA GAGTGTGGGA ATGACCGTCGGGA GGG GCGAC G
